# Discrete Multiwalled Carbon Nanotubes for Versatile Intracellular Transport of Functional Biomolecular Complexes

**DOI:** 10.1101/2023.06.06.543926

**Authors:** Kevin Castillo, Aaron Tasset, Milos Marinkovic, Aaron Foote

## Abstract

In recent years, carbon nanotubes have emerged as a potentially revolutionary material with numerous uses in biomedical applications. Compared to other nanoparticles, discrete multi-walled carbon nanotubes (dMWCNTs) have been shown to exhibit advantageous characteristics such as high surface area to volume ratio, biocompatibility, and unique chemical and physical properties. dMWCNTs can be modified to load various molecules such as proteins and nucleic acids and are capable of crossing the cell membrane, making them attractive delivery vehicles for biomolecules. To investigate this, we measured the impact of dMWCNTs on cell proliferation. Furthermore, we used electron microscopy to demonstrate that dMWCNTs enter the cytoplasm of mammalian cells via an endocytosis-like process. And lastly, we employed various *in vitro* reporter and gene assays to demonstrate dMWCNT-mediated delivery of peptides, mRNA, siRNA, and dsRNA. Our work here has helped further characterize dMWCNTs as a versatile delivery platform for biomolecular cargo.

## Introduction

The delivery of specific molecules into cells and tissues is a fundamental utility in bioengineering because it offers the potential to manipulate and control cellular behavior. Intracellular transport has been an enabling step in increasingly sophisticated biomanufacturing of products ranging from antibodies and recombinant proteins to genetically engineered cellular therapeutics [1]. More broadly, transporting molecular cargo across the plasma membrane is a prerequisite for numerous prospective therapeutic and diagnostic biotechnologies such as genome editing and intracellular imaging [2]. The cell membrane is largely impermeable to many biological molecules, making efficient trafficking into cells and tissues a significant obstacle for entire classes of molecular cargos [3]. Furthermore, once a molecule is internalized across the cell membrane, it must be transported or rely on diffusion to reach its appropriate intracellular target while minimizing degradation or adverse interference from elements in the cytosol.

Current methods of overcoming these challenges utilize either vector-based approaches, such as cationic polymers and lipid-based nanoparticle systems, or physical methods, such as electroporation and sonoporation [4–10]. Physical methods theoretically allow for direct delivery of molecules into the cytoplasm through disruption of the cellular membrane, circumventing the endocytotic process and need for endosomal escape. While efficient intracellular delivery can be achieved, these methods are often destructive in nature and require more complex setup which limits their usability to *ex vivo* and *in vitro* applications [8,11]. Nanoparticles have attracted attention for intracellular delivery due to their small size, high surface-area-to-volume ratio, and chemical tunability [9,10]. While efficient cellular uptake of nanoparticles can be achieved through chemical modification, they often suffer from increased cellular toxicity, minimizing their potential impact outside of *in vitro* applications[12,13]. Furthermore, attempts to modify these vectors to promote efficient endosomal escape and release of cargo before degradation have been largely fruitless [3,14]. While extensive work has been conducted on physical and vector-based delivery methods, identifying a versatile, systematic approach that efficiently delivers a wide variety of molecular cargo to a broad range of cell types while eliciting minimal cytotoxicity has proven elusive [15,16].

One nanoparticle species that has garnered interest for intracellular transport is carbon nanotubes (CNTs) [17,18]. These high aspect ratio, submicron cylindrical structures have unique chemical and physical properties, an exceptionally high surface area to volume ratio, and can be covalently or non-covalently modified with surfactants or biological molecules for a variety of applications [19]. CNTs exist in single-walled (SWCNT), double-walled (DWCNT) and multiwalled (MWCNT) variants and can readily bind to a wide range of biomolecules and have been shown to have great potential in drug delivery, imaging, gene editing, and other biomedical areas [20–22]. Additionally, cells appear to internalize CNTs through phagocytosis, endocytosis, and passive transport across the membrane without significantly affecting membrane integrity [22–28].

Despite their promise, investigations of carbon nanotubes have often reported results inconsistent with their posited versatile molecular loading, cell trafficking utility, and non-cytotoxic properties. The primary cause for the discrepancy between theoretical performance and experimental outcomes is a gap in understanding of how the physical and chemical properties of CNTs impact both molecular entrapment of cargo molecules and their cellular internalization. For example, studies of intracellular trafficking of MWCNTs have demonstrated that a myriad of factors, ranging from particle size and stiffness to surface chemistry and charge density, impact not only the rate of cellular uptake but the mechanism by which the particles cross the plasma membrane [29–32]. In addition to the chemical and physical attributes of the nanotubes themselves, de-bundling and dispersing of discretized MWCNTs is an essential first step for exploiting their nanoscale properties for intracellular trafficking [25].

Forming stable dispersions of discreet particles is critical for utilizing MWCNTs in biological applications. Typically, commercial grade carbon nanotubes are produced as tangled bundles of pristine carbon tubules, many microns in length. In this aggregated state, the capability of the nanotubes to serve as effective vehicles for intracellular transport is significantly reduced [29,33,34]. If CNTs are aggregated, a substantial portion of their surface area is unavailable for loading biomolecular cargo, reducing their capacity to carry relevant molecules. Additionally, aggregation limits the ability of CNTs to traffic across the cell membrane, as the relatively large bundles of particles can only be internalized via phagocytosis [23]. Lastly, aggregated CNTs have been shown to elicit toxic effects both *in vitro* and *in vivo*, limiting their application in living systems [34]. Thus, discretization of CNTs not only improves their ability to serve as effective vehicles for intracellular trafficking it also improves their biocompatibility.

In this work, we have used a proprietary process to produce discrete, surface functionalized, multiwalled CNTs (dMWCNTs), which we previously demonstrated to have minimal toxic effects *in vitro* and *in vivo* [35]. In the present investigation, we employ functionalized dMWCNTs that are ∼ 850 nm in length and ∼ 14 nm in diameter non-covalently complexed with either 1,2-distearoyl-sn-glycero-3-phosphoethanolamine-N-[amino(polyethylene glycol)-2000] (DSPE-PEG) or polyethyleneimine (PEI) for improved aqueous dispersibility and biocompatibility. We first assess the impact of dMWCNTs on the proliferation of a range of cell types at concentrations relevant for biomolecule delivery. After establishing the low *in vitro* cytotoxicity of dMWCNTs, we show that peptides, mRNA, siRNA, and synthetic dsRNA can be loaded onto carbon nanotubes and internalized by a range of cell types, including mesenchymal stem cells (MSCs), LNCaPs, T lymphocytes, and HEK cells. Once delivered into the cytoplasm, these cargoes reach their intracellular target with intact functionality and produce a measurable response. We quantified the internalization of carbon nanotubes and the effects of the corresponding cargo using various techniques, including scanning electron microscopy, fluorescence microscopy, live/dead staining, and real-time PCR. With these results, we demonstrate the potential of discrete CNTs as versatile intracellular delivery molecules. We provide critical evidence that the molecular species delivered by the particles traffic into cells and retain their bioactivity by participating in intracellular processes.

## Results

### Impact of dMWCNTs on Cell Proliferation

To demonstrate the utility of dMWCNTs as molecular delivery vehicles at the cellular level, we investigated their impact on cell proliferation. To do this, we treated Jurkat T cells, HEK cells, and MSCs with 0.001, 0.01, and 0.1 mg/ml of dMWCNTs, dispersed with DSPE-PEG(NH_2_). The change in live and dead cell numbers from day 1 to day 3 of incubation are shown in Fig 1.

**Fig. 1.**
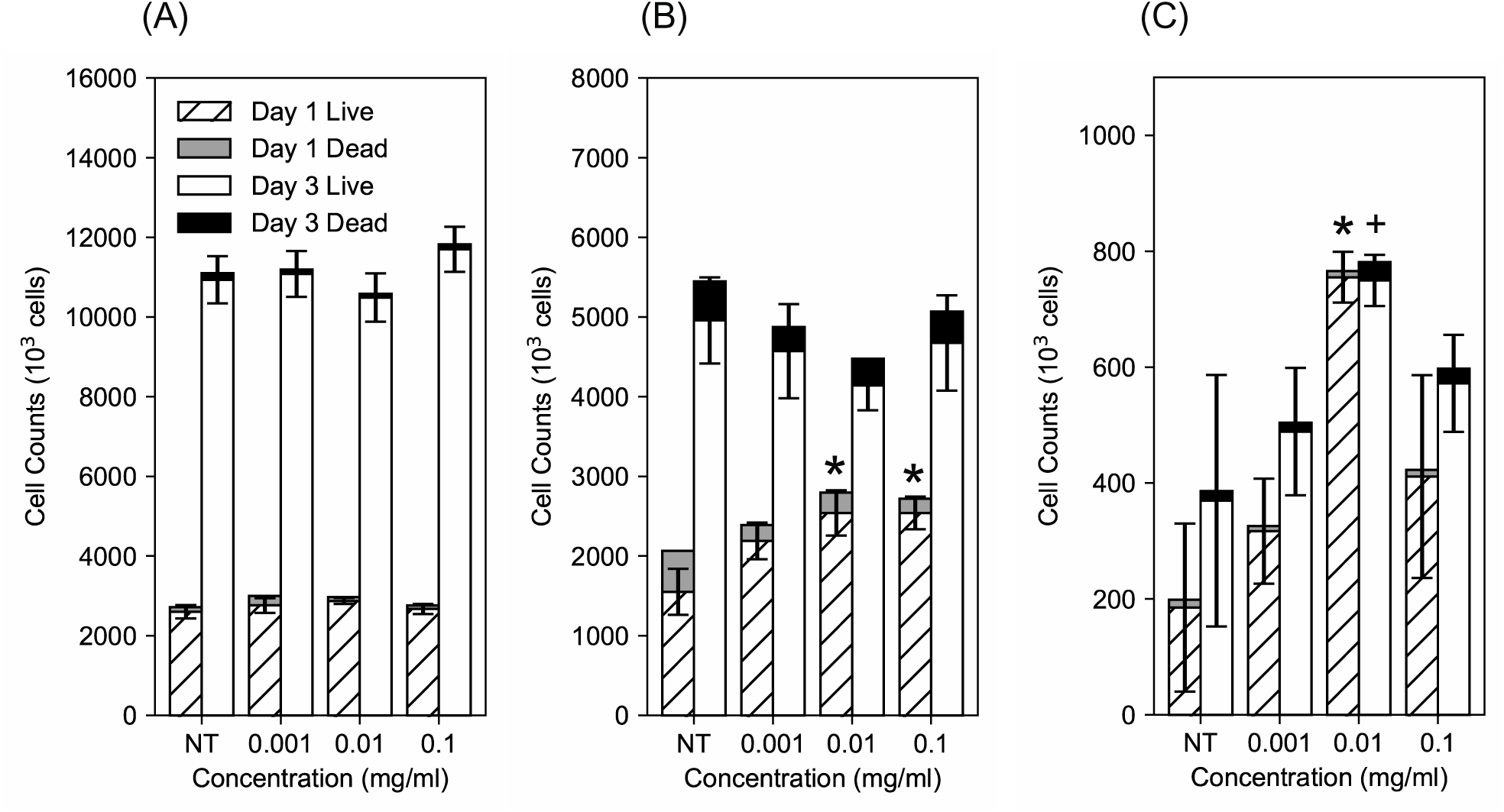
Proliferation of Jurkat T cells (A), HEK cells (B), and MSCs (C) after 1 day and 3 days of incubation with 0.001, 0.01, and 0.1 mg/ml of functionalized dMWCNTs. NT = Not Treated. Error bars represent standard error of the mean (SEM). ^✱^P< 0.05, vs. NT for Day 1. ^✚^P< 0.05, vs. NT for Day 3.

In Jurkat cells, no significant differences in cell counts were observed between the untreated group (NT) and all concentrations of dMWCNTs at the 1 and 3 day incubation time points (Fig. 1A). Additionally, the cell proliferation increase between days 1 and 3 of dMWCNT treatment is constant for Jurkat cells across all conditions tested. In HEK cells, the two highest concentrations of dMWCNTs elicited increased cell proliferation on day 1 (Fig. 1B). However, there were no significant differences between untreated cells and any of the dMWCNT-treated conditions on day 3. Finally, MSCs appeared to be affected by dMWCNTs differently than the other two cell types (Fig. 1C). Average cell densities were not significantly different after 1 day or 3 days of incubation for all dMWCNT concentrations compared to the NT group. However, the 0.01 mg/ml concentration of MGMR produced higher cell densities relative to the NT group at 1 and 3 days of culture. Interestingly, cell numbers for this treatment group did not appear to increase from day 1 to day 3. Additionally, the difference between cell densities measured on day 1 and day 3 for MSCs was not statistically significant for any condition in this study, unlike the other two cell types. Regardless of the differences between Jurkat, HEK, and MSC cell response to dMWCNTs, discrete carbon nanotubes do not appear to inhibit cell proliferation at any concentration tested.

### Internalization dMWCNTs into the Cell

To investigate the processes through which cellular internalization of dMWCNTs occurs, we incubated human bone marrow-derived (hBM)-MSCs with dMWCNTs for 72 hours. Discrete carbon nanotubes appear as black filament-like structures under TEM. In Fig. 2A. we find evidence of a single carbon nanotube (white arrow) internalized within an intracellular vesicular structure, suggesting that carbon nanotubes enters the cell membrane via an endocytosis-like mechanism.

**Fig. 2.**
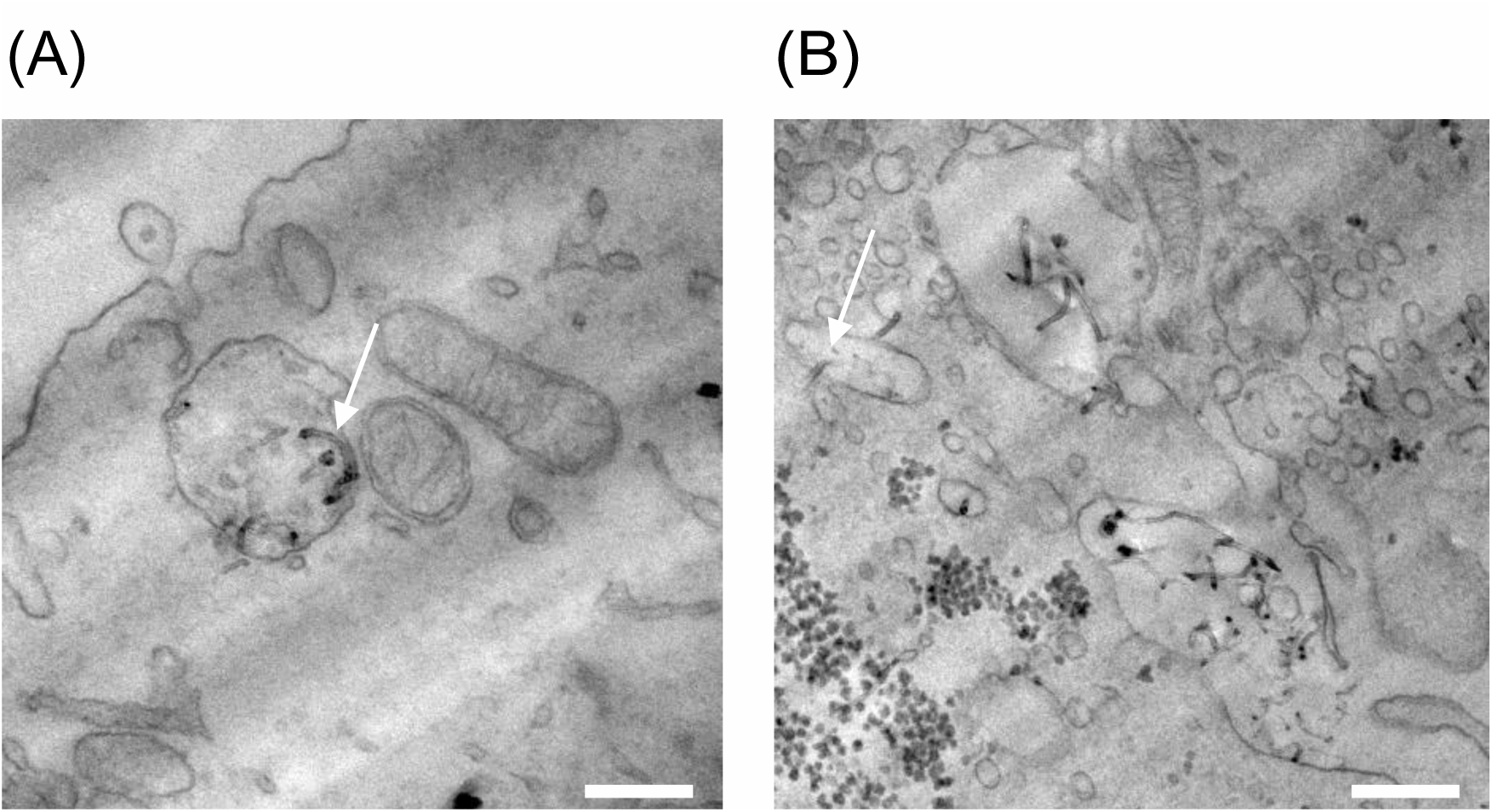
TEM Images of Cell Internalization. (**A)** Sample carbon nanotube internalized within an endosome (white arrow). **(B)** Sample carbon nanotube exiting an intracellular endosome (white arrow). Calcium deposits, appearing as round black structures, are visible in the cytoplasm. Scale bar: 200 nm.

Cytoplasmic trafficking of carbon nanotubes is strongly influenced by its discretized state. Thus, in the following experiments, we investigate dMWNCT-mediated intracellular delivery of various payloads into the cell. This demonstrates that various loaded cargo types remain functional and provides evidence of the utility of carbon nanotubes as a versatile vehicle for intracellular trafficking.

### Intracellular Delivery of KLA peptide

In this experiment, dMWCNTs were loaded with (KLAKLAK)_2_ (KLA), a membrane impermeable pro-apoptotic peptide, in a 1:2 KLA:dMWCNT mass ratio and combined with LNCaP cell culture for 48 hours. A concentration-matched control group of free KLA peptide was also prepared to account for any effects that free KLA may have on cell number. Similarly, a dMWCNT alone group was added to the study to measure any effects that the dMWCNTs may have on LNCaP cells independent of KLA delivery. Cell counts after 48 hours were performed and are shown in Fig. 3.

**Fig. 3.**
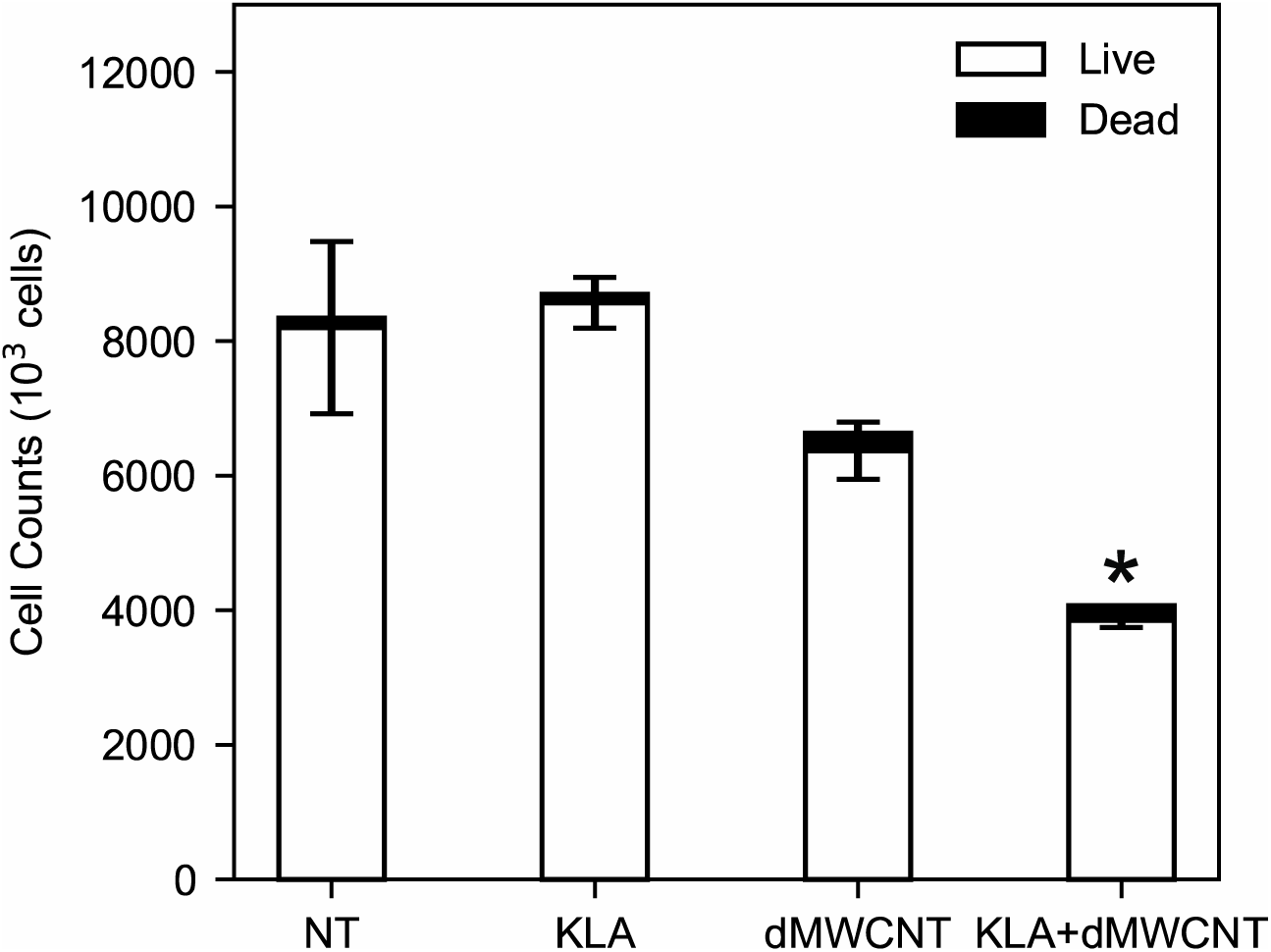
LNCaP cell number after delivery of apoptotic peptide KLA using dMWCNTs. No treatment (NT), and concentration matched controls of KLA and MWCNTs alone are included as well. Error bars represent SEM. ^✱^P< 0.05, vs. NT.

Relative to the untreated cells, the KLA alone group did not produce a significant difference in cell number after 48 hours of incubation. This observation confirms that KLA alone is membrane impermeable under the concentration and conditions used. Similarly, the dMWCNT alone group did not produce a difference in the measured cell number. However, the groups treated with dMWCNTs loaded with KLA peptide showed significant reductions in cell number, suggesting substantial apoptotic activity of the peptide-loaded on dMWCNTs. The observation that cell growth was inhibited with the combination of KLA and dMWCNTs and unaffected by either component alone) suggests that dMWCNTs facilitate intracellular trafficking and delivery of functional KLA peptides.

### Intracellular Delivery of mRNA

Next, we measured the ability of dMWCNTs to deliver genetic material into the cytoplasm for gene transfection, genetic knockdown, and T cell activation. First, we used eGFP mRNA as a model payload to quantify the level of transmembrane delivery and subsequent transient expression of mRNA delivered into cells using carbon nanotubes. We investigated mRNA internalization in two cell types: Jurkat T Lymphocytes and Monocyte-like U937 cells (Fig. 4). For both cell lines, eGFP expression was compared across three conditions: no treatment, mRNA alone, and mRNA loaded onto dMWCNTs. Cells were incubated with mRNA or mRNA bound to dMWCNTs for 72 hours at an mRNA concentration of 500 ng/ml. We used qPCR to quantify gene expression with GAPDH as the endogenous gene for both cell types and three experimental conditions.

**Fig. 4.**
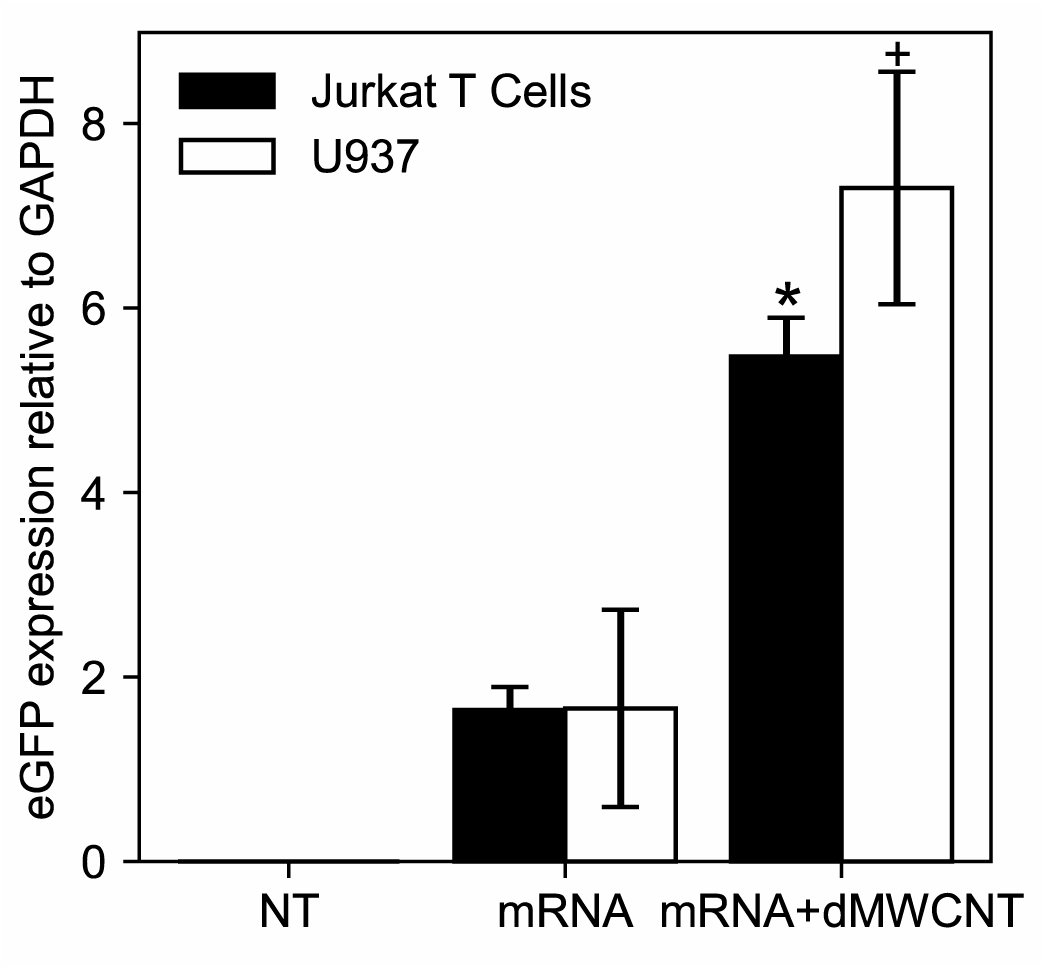
Delivery of mRNA to T-lymphocytes (Jurkat) and Monocyte-like cells (U937). eGFP expression relative to the endogenous gene GAPDH for no treatment, eGFP mRNA alone, and mRNA loaded dMWCNT groups for T-Lymphocyte Jurkats (blue) and Monocyte-like U937 (orange) cells. Error bars represent SEM. ^✱^P< 0.05, vs. NT for Jurkat. ^✚^P< 0.05, vs. NT for U937.

Incubating both cell types with free mRNA produced an approximately 1.6-fold increase in eGFP expression relative to GAPDH. However, when the same mass of mRNA is loaded onto dMWCNTs, we measured a 5.5- and 7.3-fold increase in eGFP expression for T Lymphocytes and Monocyte-like U937 cells, respectively. This increase in gene expression observed in both cell lines suggests that dMWCNTs significantly facilitate mRNA transfection, ultimately resulting in higher rates of gene expression.

### Intracellular Delivery of siRNA

To further investigate dMWCNTs as a nucleic acid delivery platform, we treated a transformed human embryonic kidney (HEK) cell line that stably expresses GFP with small interfering RNA (siRNA) with and without dMWCNTs and observed changes in GFP expression. In this experiment, successful delivery of siRNA was indicated by interference in cellular GFP expression, measured as fluorescence output. For control groups, we treated HEK cells with PBS, dMWCNTs alone, and siRNA alone to compare against the effects of siRNA combined with dMWCNTs. Using brightfield microscopy, we observed less fluorescence in cells treated with dMWCNT loaded with siRNA than in any control group (Fig.5A). To quantify GFP expression at the transcription level, we measured GFP expression of all groups relative to ACTB, normalizing to the untreated control. We found that normalized GFP expression of cells treated with unloaded dMWCNTs or siRNA only was not significantly different than untreated controls. However, cells treated with siRNA-loaded dMWCNTs exhibited a statistically significant, 0.6-fold reduction in mRNA-level expression of GFP (Fig. 5B).

**Fig. 5.**
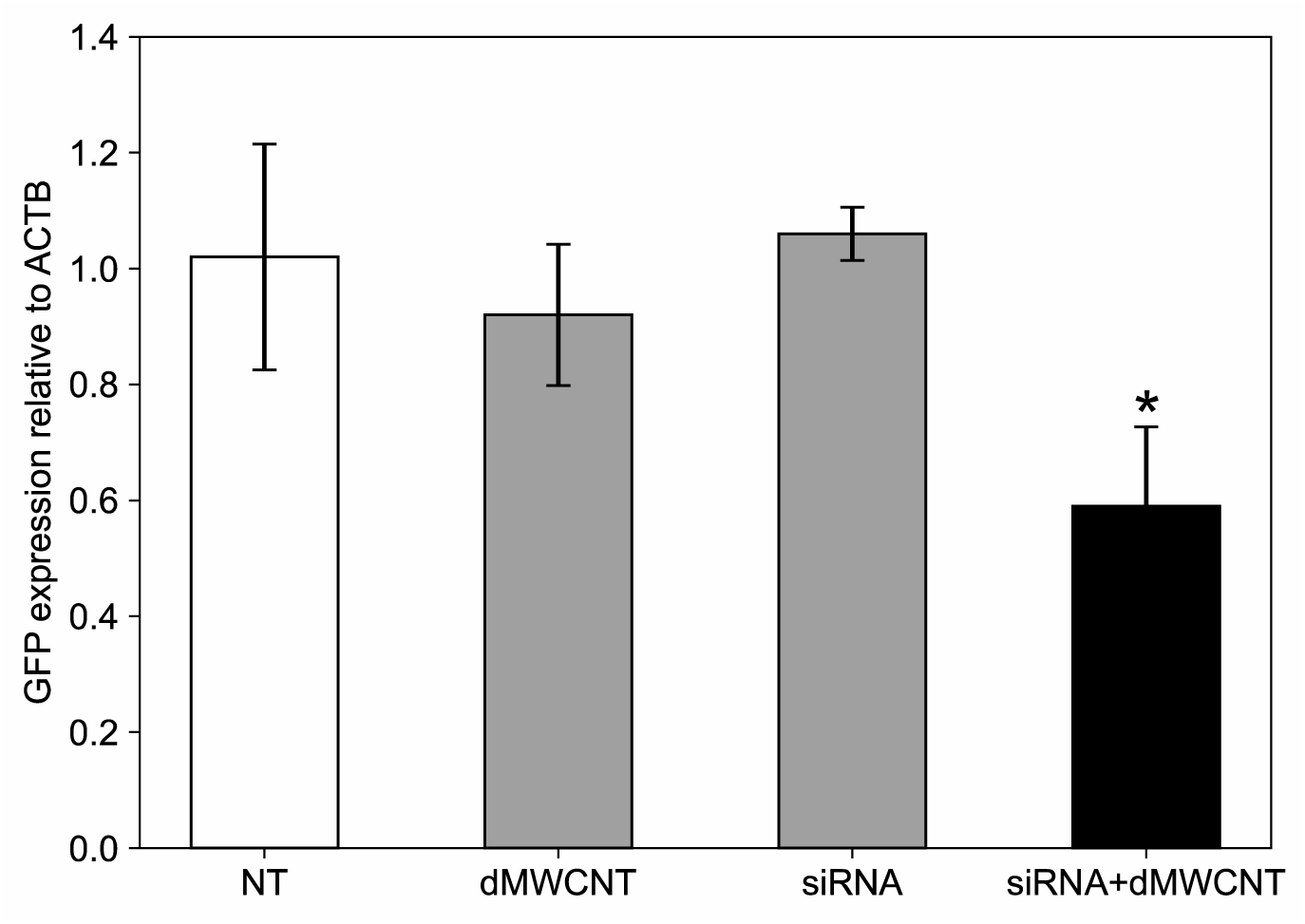
Internalization of GFP knockdown siRNA in HEK 293/GFP cells. **(A)** Fluorescence images of HEK cell cultures treated with dMWCNTs alone, GFP knockdown siRNA alone, and siRNA loaded onto dMWCNTs. **(B)** GFP expression relative to the endogenous gene ACTB in HEK 293/GFP cells treated with PBS (control). dMWCNTs only, siRNA only, and dMWCNTs loaded with siRNA. Error bars represent SEM. ^✱^P< 0.05, vs. NT.

### Intracellular Delivery of Synthetic dsRNA

Finally, we investigated the ability of dMWCNTs to deliver nucleic acids for immunoactivition. We selected the synthetic double-stranded RNA polyinosine-polycytidylic acid (Poly(I:C)), a TLR3 agonist that activates an inflammatory response when internalized within HEK cells (Alexopoulou 2001, Matsumoto 2002). Furthermore, activation by Poly(I:C) can be quantified using the HEK-Blue mTLR3 cell line, which produces the reporter secreted embryonic alkaline phosphatase (SEAP) when activated (see Methods).

Polyethyleneimine (PEI) is a cationic polymer that is commonly used to transfect cells and has high binding affinity to negatively charged nucleic acids (Pandey 2016). To improve nucleic acid loading, we physisorbed 270 kDa branched PEI to the nanotube surface, creating a modified dMWCNT. However, to distinguish between the delivery capability of PEI alone and dMWCNTs, we incorporated a concentration-matched control group of Poly(I:C) loaded onto PEI alone. In this way, relative HEK-Blue mTLR3 activation was quantified upon treatment with free Poly(I:C), Poly(I:C) complexed with PEI, and Poly(I:C) loaded onto carbon nanotubes physisorbed to PEI (Fig. 6).

**Fig. 6.**
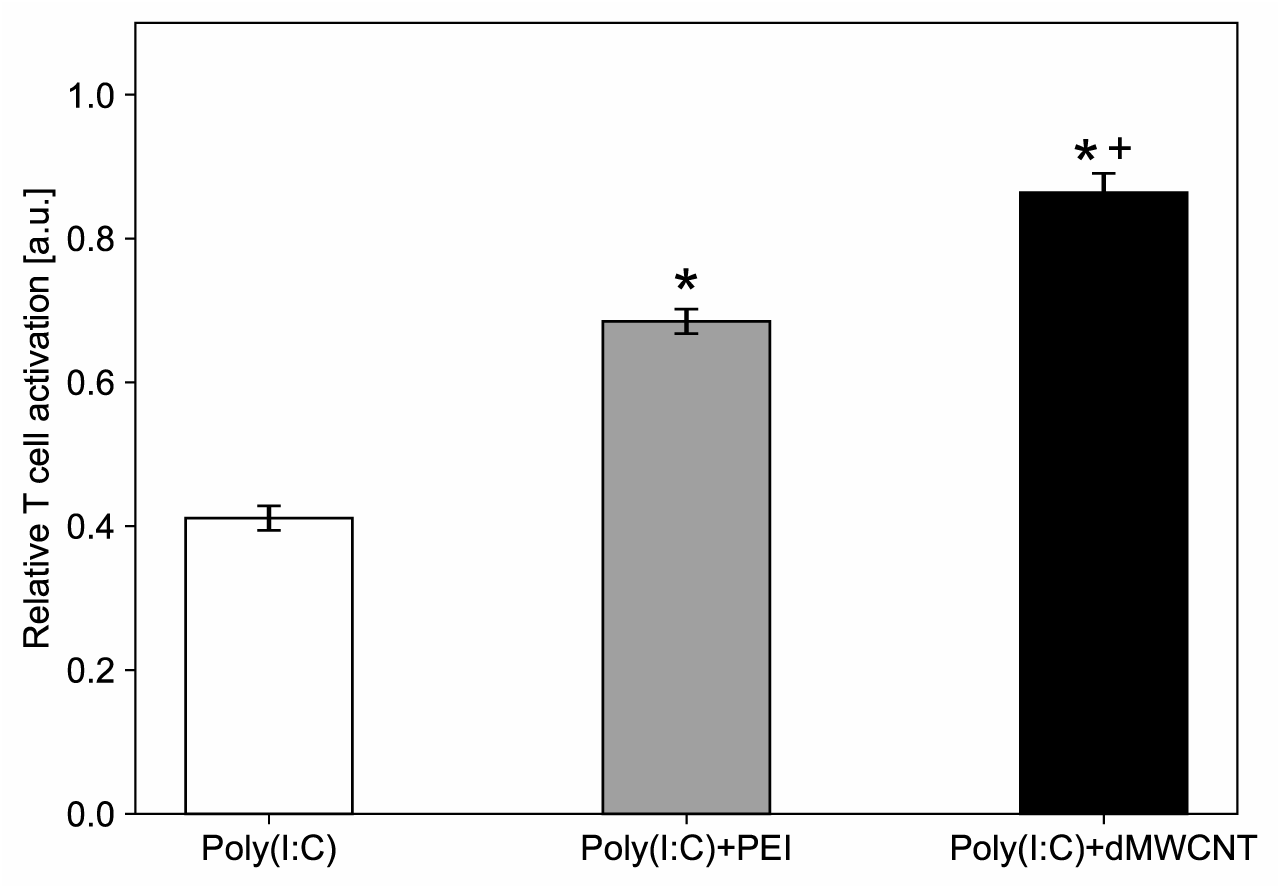
Intracellular delivery of Poly(I:C) to HEK-blue cells as measured by relative T cell activation. Cells were treated with Poly(I:C) alone (control), Poly(I:C) loaded onto PEI, and Poly(I:C) loaded onto dMWCNTs physisorbed with PEI to the carbon nanotube surface. Error bars represent SEM. ^✱^P< 0.05, vs. Poly(I:C). ^✚^P< 0.05, dMWCNT vs. PEI.

Comparing the Poly(I:C) alone control and the Poly(I:C) with PEI alone group, we measured a significant increase in HEK cell activation when Poly(I:C) is bound to PEI, in agreement with the literature. However, we observed the greatest degree of T cell activation from the Poly(I:C) loaded onto dMWCNTs physisorbed to PEI. This group produced approximately an 100% improvement from the Poly(I:C) alone and a ∼20% improvement from the PEI complexed Poly(I:C) group.

## Discussion

Discretizing carbon nanotubes and preventing their aggregation is essential for their use as biocompatible delivery vehicles [29,33]. While aggregated CNTs strongly decrease cell and tissue viability [34], discrete MWCNTs do not demonstrate cytotoxic effects [35]. Furthermore, discrete tubes internalize into cells more efficiently and have more available surface area for biomolecule loading.

This work demonstrates how dMWCNTs act as versatile intracellular delivery vehicles that do not moderate cell proliferation at concentrations practical for intracellular trafficking. Utilizing TEM imaging, we presented evidence that the dMWCNTs used in this study were de-bundled and internalized into mesenchymal stem cells and, critically, can escape from intracellular vesicles in order to delivery their biomolecular cargo into the cytoplasm (Fig. 2). Though the exact mechanisms are not fully understood, the micrographs in Figure 2 suggest that internalization appears to occur through a vesicle-mediated process, followed by vesicular escape and release of the dMWCNTs into the cytoplasm. Importantly, the nanotubes used in our study retained their discreet state and did not appear to interact with the intracellular cytoplasmic mineralized calcium produced by differentiating hBM-MSCs [36,37]. Biomolecular payloads including the peptide KLA, mRNA, siRNA, and dsRNA were delivered in LNCaP, Jurkat, U937, and HEK cells using dMWCNTs. Furthermore, the functionality of delivered biomolecules was preserved in each case tested.

In order to serve as an effective and versatile delivery vehicle for biomolecules, dMWCNTs carrying a molecular payload must not only enter the cytoplasm but release their cargo within a reasonable time scale without altering its biological functionality. In this regard, a potential advantage of the nanotubes prepared in our study is that their dispersed and discretized state, high surface-to-volume ratio and variety of charge densities in its polymeric/lipid coating (i.e. DSPE-PEG), supported numerous types of electrostatic interactions (i.e. hydrogen bonding, and ionic, van der Waals, and hydrophobic interactions) useful for biomolecular loading. In addition, because each of the cargo molecules we tested complexed with dMWCNTs through non-covalent chemical bonding, it is likely that they were eventually released from the surface of the particles by simple diffusion and/or replacement by other chemical species inside the cell. While this approach does not offer the same degree of control over biomolecular payload delivery as covalent interactions, it does not require structurally modifying the cargo molecules or breaking chemical bonds in order to release into the cytoplasm.

In order to evaluate this approach, the pro-apoptotic peptide KLA was selected as a representative cargo molecule to evaluate this capability. KLA is membrane impermeable but toxic to mitochondrial membranes if internalized into LNCaP cells, resulting in apoptosis [38–41] . As its apoptotic mechanism of action relies on its ability to disrupt the cellular mitochondrial membrane, its effectiveness is susceptible to conformational changes of the peptide [42]. For these reasons, KLA serves as a helpful test payload to validate that physisorption on the carbon nanotube surface does not cause changes that impair its functionality.

In the case of KLA delivery, we observed a slight decrease in cell number in the dMWCNT alone group when compared to the no treatment condition (Fig. 3). In this case, the concentration of dMWCNT used for KLA loading was 0.138 mg/ml, a relatively high value when compared to the proliferation studies performed in Fig. 1 (0.001 mg/ml, 0.01 mg/ml, 0.1 mg/ml). The high concentration was needed in order to ensure the binding of KLA to some amount of dMWCNTs in solution, as we found that the binding affinity of KLA to dMWCNT to be relatively low. This affinity could be improved in future studies by further modifying the surface of the dMWCNT. However, the physisorption and desorption of bimolecular payloads requires fine tuning of binding affinities, as a biomolecular payload that is too strongly adhered to the nanotubes surface may not be able to perform its desired function inside of the cell.

Similarly, binding affinity may also have been an important factor in both experiments involving intracellular delivery of nucleic acids. The particles were shown to significantly improve transfection efficiency relative to uptake of free mRNA (Fig. 4). Similarly, we observed that siRNA alone did not reduce GFP expression, however loaded dMWCNTs were shown to be an effective vehicle for siRNA transfection into the cytoplasm (Fig. 5). Although the combination of siRNA and dMWCNTs markedly improved cellular uptake and resulted in approximately 40% knockdown of gene expression, it is likely that transfection effectiveness could have been improved by further optimization of siRNA interaction with the dMWCNT surface chemistry. In particular, a higher degree of control over off-loading kinetics may improve the release profile of the molecular payload and increase transfection efficiency.

Another aspect of using dMWCNTs for intracellular biomolecule delivery that is worth further investigation is the protective effect that nanotubes may have on molecular degradation. Nucleic acids, such as the mRNA and siRNA delivered in Fig. 4 and Fig. 5, were shown to be delivered intracellularly and have a stronger effect when loaded onto dMWCNTs than when incubated with cells without any delivery vehicle. Though this could be partially or entirely explained by the dMWCNTs ability to cross the cell membrane, it is a strong possibility that carbon nanotube loading prevents nucleases from damaging the bound nucleic acids. Optimizing this protective effect could lead to the use of dMWCNTs as transfection agents for cell and gene therapy applications.

Finally, the Poly(I:C) used in this study is a comparatively large nucleic acid (1.5-8 kbp) in comparison to the mRNA and siRNA tested earlier in this work (Fig. 6). Poly(I:C) is an immunostimulant, which has been studied for decades as a vaccine adjuvant [43]. While a previous study used single-walled carbon nanotubes (SWCNTs) for intracellular delivery of poly I:C [44], to our knowledge, our study is the first to describe to use of dMWCNTs for this purpose. The use of dMWCNTs as vehicles for adjuvant delivery, or as adjuvant materials in their own right, could be an attractive intracellular delivery modality and warrants further study. Furthermore, the increase in cell activation observed when Poly(I:C) is loaded onto dMWCNT physisorbed to PEI compared to Poly(I:C) complexed with PEI alone suggests that the nanotube provides a benefit independent of PEI activity. It is possible that carbon nanotubes physisorbed with other transfection or drug delivery agents could provide a similar benefit, opening up additional avenues of biomedical applications of dMWCNTs.

## Materials and Methods

### CNT Functionalization and Discretization

The MWCNTs used in this work are oxidized and de-bundled using a process previously described (Falank 2019). The resulting wetcake of functionalized carbon nanotubes was then dispersed via water bath sonication in DSPE-PEG (MW = 2000, Laysan Bio) or by covalently binding mPEG (MW=5000, Sigma Aldrich) followed by further dispersal in DSPE-PEG, depending on the experiment. Dispersal of oxidized MWCNTs in PEI (270 kDa, Sigma Aldrich) was also performed using a water bath sonicator and was used to load and deliver Poly(I:C) intracellularly. The discretized nature of each formulation of dMWCNTs was confirmed via brightfield microscopy.

### Cell Internalization and TEM Imaging

MSCs were cultured in GM containing 0.001 mg/mL dMWCNTs for 72 hours. Before fixation, the cells were washed with PBS. Tissues were dissected in PBS and exchanged into 0.1M Sodium Cacodylate buffer prior to fixation in 4% glutaraldehyde with 3% paraformaldehyde in the same, and then staining with 1% osmium tetroxide in the same, followed by 1% uranyl acetate in water. After dehydration through graded alcohols and embedding in resin (Hard Plus Epon 812, EMS), thin sections were cut on a Leica UC7 ultramicrotome and mounted to grids. Images were captured on a Tecnai Spirit TEM at 80kV.

### Proliferation/cytotoxicity assay

Cell lines were seeded in a 12-well flat bottom plate at a concentration of 1.0x10^5^ cells/mL per well. Cells were then incubated for 24 hours in growth media (GM), consisting of α-MEM (Life Technologies, Grand Island, NY) containing glutamine (2 mM), penicillin/streptomycin, and 15% fetal bovine serum (FBS) (Atlanta Biologicals, Lawrenceville, GA). After 24 hours, cells were treated with respective experimental groups and controls (0.1 mg/ml, 0.01 mg/ml, 0.001 mg/ml dMWCNTs). The dMWCNT formulation used in this study consisted of carbon nanotubes dispersed in DSPE-PEG (MW = 2000, Laysan Bio). After treatments, cells were then collected from the plate at the 24- and 72-hour time points and then stained with trypan blue for live/dead counting via Thermo Fishers Countess II.

### KLA Delivery Study

dMWCNTs were loaded with KLA in a 1:2 KLA:dMWCNT mass ratio (0.069 mg/ml KLA with 0.138 mg/ml dMWCNTs) and combined with LNCaP cell culture for 48-hours. A concentration-matched control group of free KLA peptide was also prepared. A dMWCNT alone group was added to the study at a concentration of 0.138 mg/ml. The dMWCNT formulation used in this study consisted of carbon nanotubes dispersed in DSPE-PEG (MW = 2000, Laysan Bio). Cellular viability was then assessed via trypan blue staining.

### mRNA Delivery Study

eGFp mRNA (StemMACS eGFP, Miltenyi Biotec) was loaded onto dMWCNTs in a 1:500 mRNA:dMWCNT mass ratio. Groups containing mRNA were treated with 500 ng of eGFP coding mRNA. The dMWCNT formulation used here consisted of MWCNTs covalently bound to mPEG (MW = 5000 Da, Sigma Aldritch) and further dispersed in DSPE-

PEG(NH_2_) (MW = 2000 Da, Laysan Bio). RNA was isolated from tissue samples using Trizol reagent (Invitrogen) following the manufacturer instructions. Single stranded cDNA was synthesized from the isolated RNA with High-Capacity cDNA Reverse Transcription Kit from Applied biosystems. The expression of eGFP was analyzed via qRT-PCR using Taqman by Quantstudio 3 Real-Time PCR system.

### siRNA Delivery Study

Adherent HEK-293/GFP cells (Cell Biolabs, Cat. No. AKR-200) were grown in RPMI-1640 (10% FBS, 1% Anti-anti) to a confluency of 90% at 37C with 5% CO2 prior to harvesting for study. The harvested cells were then diluted to a concentration of 3.0x10^6^ cells/mL in RPMI-1640. Diluted cells were then seeded in a flat bottom 12-well plate at a final cell density of 1.0x10^5^ cells per well. After 24-hours, growth media was removed and replaced with Opti-MEM. Cells were then treated with experimental conditions PBS, siRNA alone (Stealth RNAi siRNA GFP, Thermo Fisher), siRNA-dMWCNT and incubated for 72-hours. 50 µM of siRNA was used for each treatment of siRNA containing groups, and 50 µM of siRNA was loaded onto 5 mM dMWCNTs in a 1:100 ratio for dMWCNT containing groups. The dMWCNT formulation used here consisted of MWCNTs covalently bound to mPEG (MW = 5000 Da, Sigma Aldritch) and further dispersed in DSPE-PEG(NH_2_) (MW = 2000 Da, Laysan Bio). Cells were then imaged and harvested at the 72-hour timepoint for real-time PCR analysis (ThermoFisher, QuantStudio 3).

### T-Cell Activation via PolyI:C

Human T Lymphocytes; Jurkat-Dual (Invivogen), cells were grown to a confluency of 70-80%, as per manufacturer instructions. Suspended cells were then harvested and diluted to a concentration of 5.0x10^5^ cells/mL in a selection medium (IMDM supplemented with 10% FBA, Penicillin-streptomycin). Cells were seeded in a flat bottom 96-well plate at a final concentration of 3.5x10^5^ cells/well and then treated with experimental conditions, Poly I:C (Invivogen) alone, PEI loaded with Poly(I:C), and dMWCNTs dispersed in PEI loaded with Poly(I:C). The concentration of PEI selected for loading Poly(I:C) was based on the mass of PEI physisorbed onto the surface of the corresponding dMWCNT dispersion. This mass was measured using thermogravimetric analysis as previously described (Falank 2019). Cells were incubated for 18hr at 37 °C with 5% CO_2_. Cells were then pelleted (150 RCF, 5 min) and SEAP enriched supernatant was then utilized for detection media (QUANTI-Blue, Invivogen) activation.

### Statistical Analysis

Student’s t-test was used to determine significance in all pairwise statistical comparisons; P<0.05 considered significant for all comparisons. Error bars in replicate data sets were represent standard error of the mean (SEM). The multi-way statistical comparison in Figure 6 was performed using one-way ANOVA with Tukey’s post-hoc test. At a minimum, experimental replicates were performed in triplicate and each experiment was repeated a minimum of 3 times.

## Notes

### Competing Interest Statement

Dr. Aaron Foote and Kevin Castillo are current employees of BioPact, LLC. Dr. Milos Marinkovic and Aaron Tasset are former employees of BioPact, LLC.

